# Dorsal CA1 silencing degrades tactile short-term memory

**DOI:** 10.1101/2024.03.01.582962

**Authors:** Shirly Someck, Netta Katz, Amir Levi, Eran Stark

**Affiliations:** Sagol School of Neuroscience, Tel Aviv University, Tel Aviv 6997801, Israel; Department of Physiology and Pharmacology, Faculty of Medicine, Tel Aviv University, Tel Aviv 6997801, Israel; Sagol Department of Neurobiology, Haifa University, Haifa 3103301, Israel

## Abstract

Previous studies in rodents showed that the hippocampus is involved in spatial short-term memory (STM), but hippocampal necessity for maintaining short-term sensory memories is unknown. Here, we develop tactile discrimination and STM tasks for freely-moving mice. Subjects learn to discriminate between textures after four shaping sessions and a single post-shaping session, and learn the STM task within a dozen sessions. Transient closed-loop silencing of dorsal hippocampal region CA1 during memory maintenance degrades task performance, compared to interleaved control blocks. Thus, uninterrupted hippocampal activity is required for acting upon tactile information maintained in STM. The findings suggest that the role of the hippocampus extends beyond spatial navigation, encoding memories, and long-term consolidation of experiences.

## Introduction

Keeping a memory and acting upon it are essential functions for living organisms. Memory is categorized in multiple manners depending on duration (e.g., long term memory and STM), awareness (explicit and implicit), and functionality (working memory [WM] and reference memory^1^). At short time spans, the difference between explicit STM and WM is functional: WM is useful for carrying out an action^2^, whereas STM is fully defined by storage duration and capacity^1^. Operationally, WM can be segmented into sequential phases, thought to rely on different neuronal processes: encoding, maintenance, and retrieval^3^. When the duration is short, typically up to about twenty seconds^4^, the maintenance phase of WM is considered STM.

The hippocampal formation is involved in processing navigation and higher-order spatial information^5–8^. Furthermore, the hippocampus is involved in spatial WM^9–16^ and in consolidating and retrieving long-term memories^1,17–19^. Because the involvement of the hippocampus in spatial WM may be explained by spatial or navigational components, it has been unclear whether the hippocampus is necessary for non-spatial STM.

In rodents, most previously-employed STM tasks were spatial WM tasks, including delayed alternation^11,13,20–22^ and delayed (non)match-to-sample^16,23,24^ tasks. Non-spatial STM may involve brain circuits distinct from spatial STM. Non-spatial STM tasks for rodents typically employed visual^25–30^, olfactory^31–34^, auditory^35–41^, or tactile^40–45^ sensory cues. However, all previous non-spatial STM tasks for mice employed fixation of the head^34,38,43^ or the entire body^33^. Spatial STM tasks are usually faster to learn and yield more trials compared with non-spatial tasks (**Table S1**), possibly because the spatial tasks exploit the exploratory nature of rodents. Thus, while there are multiple spatial STM tasks for rodents and various sensory STM tasks for freely-moving rats and for head-fixed mice, there are no sensory STM tasks for freely-moving mice.

## Results

### Mice learn to discriminate between textures within five sessions

In comparison to odor and taste, tactile stimuli can be easily limited in duration. In comparison to vision, palpable stimuli are ecological and organic to mice. Indeed, tactile discrimination (TD) has been widely studied in rodents (**Table S2**) and tactile tasks are learned rapidly^46–48^. Many previously-employed TD tasks included object localization and texture discrimination tasks^49–54^ while the rodent was head-fixed. Here, we designed an automated apparatus that enables tactile discrimination and STM tasks in freely-moving mice. The apparatus is a figure-8 maze with two pairs of texture wheels at the beginning and end of the central arm (**Fig. 1A**). Tactile stimuli were presented by two pairs of motorized wheels, equipped with sandpaper of varying grits on four facets and a single smooth (null) facet. Stimulus-response contingency was fixed, and in all sessions P60 (coarse) textures were associated with leftward runs, whereas P320 (extra fine) textures were associated with rightward runs (**Fig. S1A**). The apparatus supports at least two distinct cognitive tasks: a TD task and a tactile delayed-response STM task.

**Figure 1.**
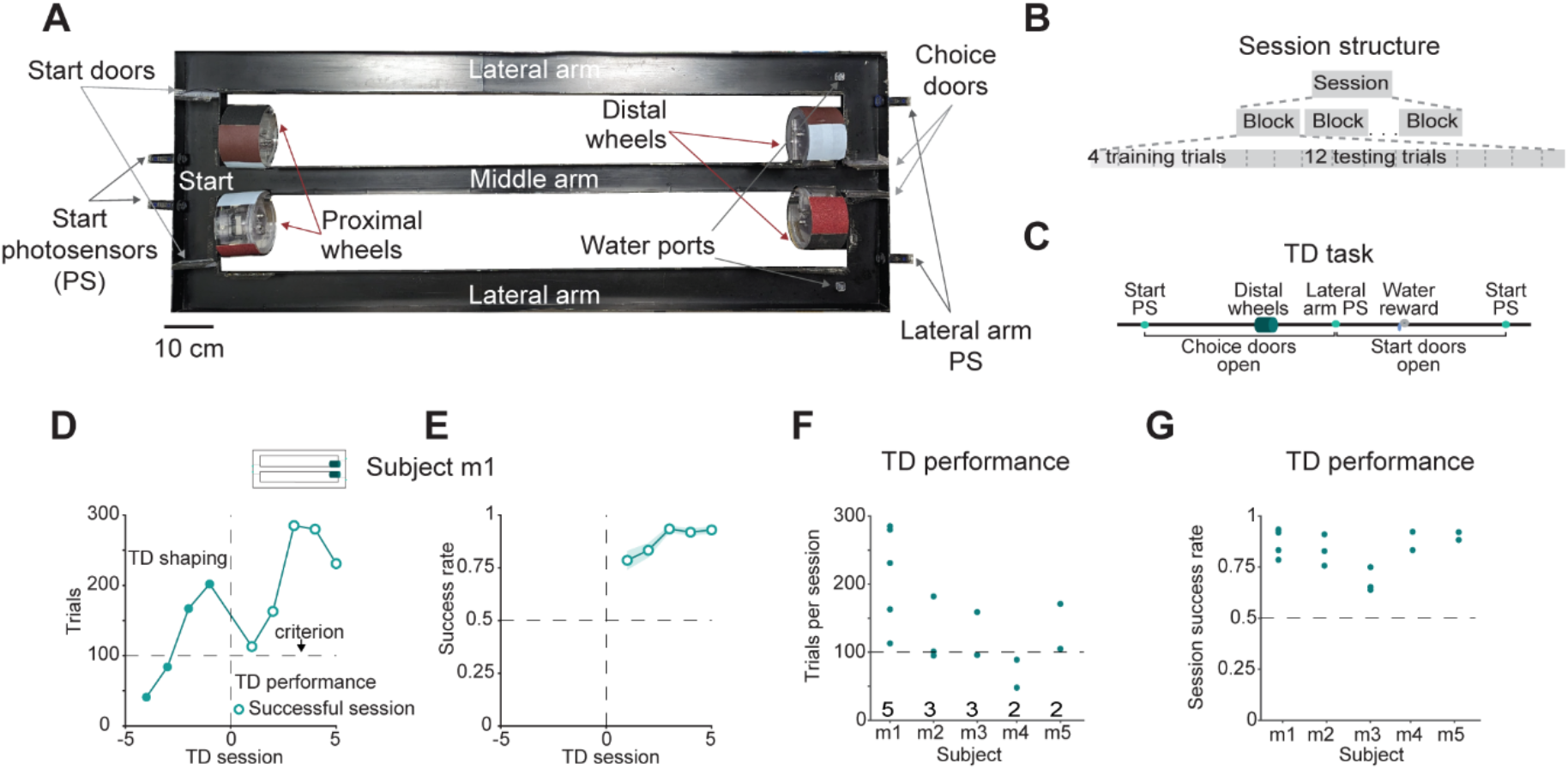
Mice learn to discriminate between textures within five sessions. (**A**) Figure-8 maze with two pairs of motorized texture wheels situated 120 cm apart. Each pair of wheels consists of four facets with sandpaper grits of P60 (coarsest), P150, P240, and P320 (finest); and a single smooth (null) facet. (**B**) Every session is divided into blocks, each composed of four training and 12 testing trials. In training trials, only the correct choice door opens, forcing the animal to choose the rewarded side. In testing trials, both doors open. (**C**) Timeline for a TD trial. (**D**) Number of trials per TD session in subject m1. Counts include training and testing trials. Here and in **F**, horizontal line, hundred-trials “shaping” criterion. Here and in **E**, successful sessions, p<0.05 in a Binomial test, comparing to chance level of 0.5; vertical line, transition between the shaping and performance phases. (**E**) Success rates (testing trials only) for the same subject as in **D**. Error band, SEM. Here and in **G**, horizontal line, chance level. (**F**) Total number of trials per TD session during the performance phase of subjects m1-m5. The number of sessions per mouse is denoted below every dot plot. (**G**) Success rates during the TD performance phase.

For either task, the mice (**Table 1**) went through a training and testing process that included up to three phases: shaping, learning, and performance (**Fig. S1BC**). The “shaping” phase continued until performing at least 100 training trials in a single session. During a training trial, the subject was presented with the texture stimulus, but the correct choice was enforced by opening only one of the choice doors at the T-junction (**Fig. 1A**). Shaped mice underwent a series of learning sessions that also included testing trials, in which both choice doors were open. To facilitate rapid learning and minimize frustration, every learning or performance session was divided into blocks (**Fig. 1B**). Each block consisted of four training trials followed by 12 testing trials. During the TD task, only the distal wheels were active, and the proximal wheels were stationary with the null (smooth) facet facing the middle arm (**Fig. 1C**). We state that the mouse has learned the task if two consecutive sessions were successful (p<0.05 on a Binomial test for all same-session testing trials), and define the time of learning as the first of the two sessions.

**Table 1.** Behavioral performance and silencing effect.

| Animal ID | Animal number | Sex | Strain | Age<sup>1</sup> [week] | Weight<sup>1</sup> [g] | TD performance<sup>2</sup> | STM performance<sup>2</sup> | Silencing effect<sup>3</sup> |
| --- | --- | --- | --- | --- | --- | --- | --- | --- |
| mA108 | m1 | F | HYB | 18 | 24.3 | 0.92 (n=5) | - | - |
| mA271 | m2 | F | HYB | 6 | 20.5 | 0.83 (n=3) | 0.66 (n=9) | - |
| mA303 | m3 | F | HYB | 78 | 28.1 | 0.65 (n=3) | - | - |
| mA355 | m4 | F | HYB | 6 | 18.5 | 0.88 (n=2) | 0.58 (n=10) | 16.7% (n=2) |
| mA356 | m5 | F | HYB | 6 | 18.5 | 0.9 (n=2) | 0.66 (n=13) | - |
| mP79 | m6 | M | C57 | 25 | 32.8 | - | 0.6 (n=16) | - |
| mA335 | m7 | M | HYB | 7 | 28.5 | - | 0.67 (n=13) | 7.7% (n=10) |
| mA352 | m8 | F | HYB | 18 | 22.2 | - | 0.72 (n=3) | 16.7% (n=3) |
| mA353 | m9 | F | HYB | 18 | 22.6 | - | 0.66 (n=36) | 16.7% (n=10) |
| **Median [IQR]** |  |  |  |  |  | **0.83 [0.73 0.88]** | **0.63 [0.59 0.68]** | **16.7% [0% 16.7%]** |

<sup>1</sup>Before the first behavioral session.
<sup>2</sup>Median success rate (number of sessions).
<sup>3</sup>Silencing effect (number of paired blocks).

The shortest training duration reported for a rodent TD task is three days^55^, in a task based on novel object recognition that did not involve individual trials. Previous trial-based TD tasks in rodents required weeks of training^26,50,51,56–58^ (**Table S2**). We found that in the present TD task, both habituation and learning occurred during the shaping phase. For example, subject m1 performed an increasing number of trials during consecutive shaping sessions (**Fig. 1D**). After crossing the hundred-trial shaping criterion, learning sessions were initiated. However, during the first session with testing trials, 66/84 (79%) of the testing trials were correct and performance was already above chance (p<0.001, Binomial test; **Fig. 1E**). Success rate increased during the next session and stabilized at 92-94%. Therefore, while asymptotic performance required more sessions than the shaping phase, m1 learned the TD task by the end of the first post-shaping session. Similar results were observed for all n=5 subjects, for whom a median [range] of 4 [4,6] shaping sessions was required to reach the hundred-trial shaping criterion (**Table S3**). All mice trained on the TD task performed successfully during the first post-shaping session (**Fig. S1D**). Thus, all mice learned to discriminate textures by the end of the first post-shaping session, within a median of five sessions.

Every mouse carried out at least two post-shaping sessions to quantify and optimize performance. Because performance during the first post-shaping session was already successful, we grouped together all post-shaping sessions. While the number of trials depended on the chronological day of the training (rank correlation, 0.66; n=15 sessions; p=0.0091, permutation test; **Fig. S1E**), all (15/15; 100%) performance sessions were successful (**Fig. S2A-E**). Over n=15 sessions in the five mice, the median [interquartile range, IQR] number of trials per session was 113 [95 182] (**Fig. 1F**), and the success rate was 0.83 [0.73 0.88] (**Fig. 1G**). Thus, after rapid learning of the TD task, performance is consistently successful.

### Mice learn the STM task within a dozen sessions

Most previously-described STM tasks for freely-moving mice were deliberately based on spatial STM, aiming to exploit the exploratory nature of the subjects and requiring a minimum of three sessions to learn^16,23,59–61^ (**Table S1**). Non-spatial STM tasks for freely-moving rodents were developed for rats, requiring a minimum of nine sessions^35,39^, but none have been reported for mice. Here, we combined whisking (sample) and running (maintenance) in the same task, yielding an organic maintenance period of a tactile STM task.

Before being exposed to the STM task (**Fig. 2A**), subject m5 was pretrained on the TD task, carrying out five shaping sessions and two successful performance sessions. After the seven TD sessions, m5 was shifted over to the STM task, using the exact same stimulus-response contingency but at the proximal wheels (**Fig. 2B**, pink), requiring memory maintenance throughout the middle arm run. In contrast to TD training which required only one post-shaping day (**Table S3**), training on the STM task required a longer learning phase (**Fig. 2C**). Following one shaping session and four learning sessions, m5 managed to learn the STM task. During the performance phase, m5 performed 11/14 (79%) successful sessions with a median [IQR] success rate of 0.63 [0.57 0.66].

**Figure 2.**
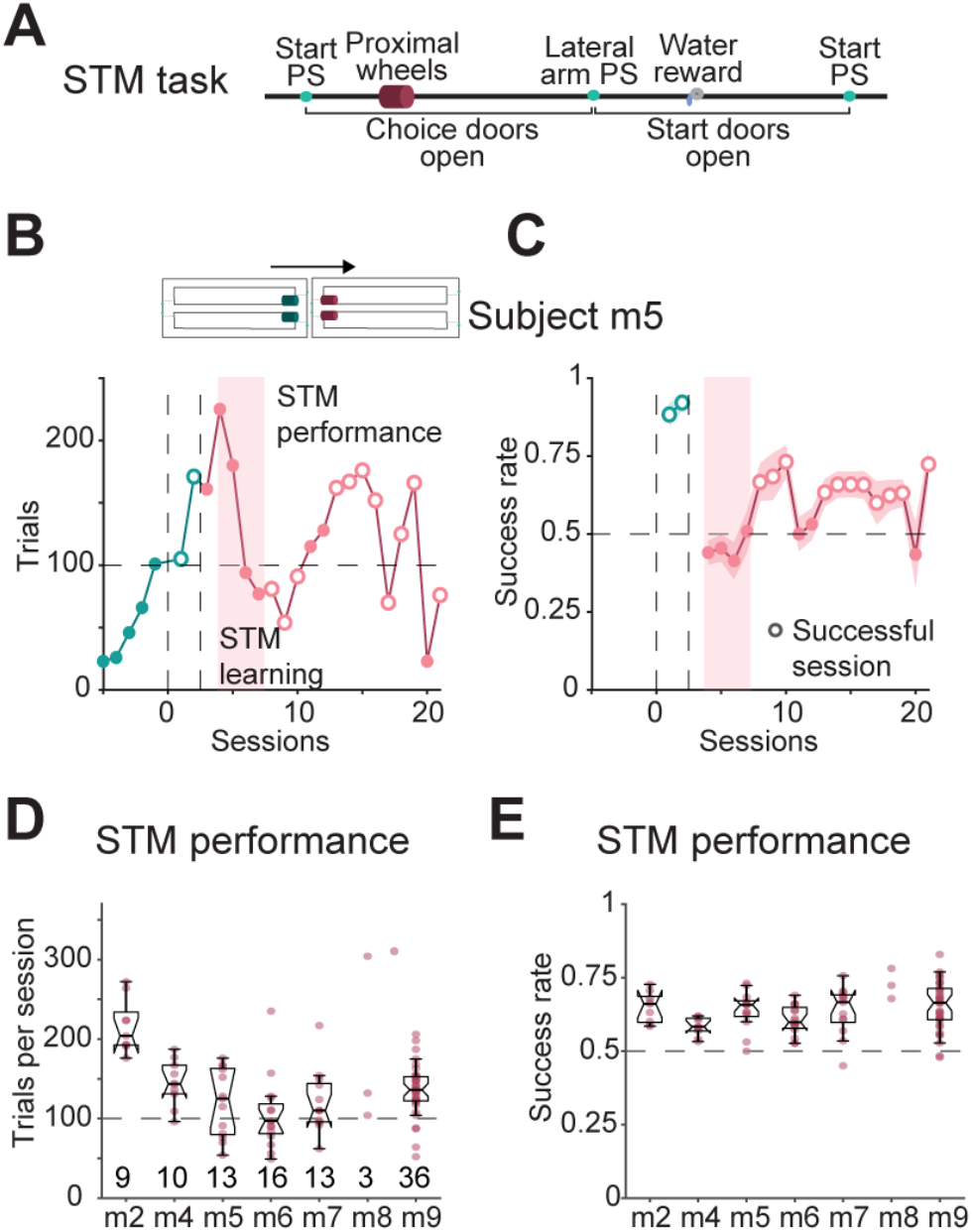
Mice learn the STM task within a dozen sessions. (**A**) Timeline for an STM trial. (**B**) Number of trials per session in subject m5 on the STM task after pretraining on the TD task (green). After STM shaping (pink background), STM training continues until the task is learned, defined as two successful STM sessions performed consecutively. Here and in **C**, all conventions are the same as in **Fig. 1D**. The second vertical line separates the TD and STM tasks. (**C**) Success rates for subject m5. (**D**) Total number of trials per session during STM performance for all seven subjects. Here and in **E**, silencing sessions (**Fig. 3**) are not included; every box plot shows median and IQR; whiskers extend for 1.5 times the IQR in every direction; and dots indicate individual sessions. The number of sessions per mouse is denoted below every box plot. (**E**) Success rates of the mice on the STM task during the performance phase.

Considering all n=7 subjects trained on the STM task, the mice spent a median [IQR] of 828 [194 2726] trials (**Fig. S1K**; **Table S3**) and 5 [1 19] sessions in the STM learning phase (**Fig. S1L**). To assess performance on the STM task regardless of training conditions (pretrained/naïve, dark/light; **Fig. S1MN, Fig. S2**), we pooled together the results of all performance sessions. 87/100 (87%) of the sessions were successful (n=7 mice). The median [IQR] number of trials per performance session was 132 [98 157] (**Fig. 2D**), and success rate was 0.63 [0.59 0.68] (**Fig. 2E**). Thus, all mice trained on the STM task succeed in learning the task, perform the task with high trial yield, and maintain above-chance performance after learning.

### Bilateral silencing of dorsal CA1 during memory maintenance degrades success

Previous studies correlated behavior with region-level neuronal activity using rapid and temporary optogenetic silencing of neuronal activity during behavior^16,24,43,45,62^, as opposed to lesions^10,61^ or pharmacology^12,36,37,63^. To determine involvement in tactile STM, we transiently silenced the dorsal hippocampus during the memory phase of the task. We injected a viral vector bilaterally into hippocampal region CA1 of four mice trained on the STM task (**Table 1**), inducing the expression of the red-light activated neuronal silencer Jaws^64^ under the CaMKII promoter (**Fig. 3AB**). We implanted optical fibers coupled to miniature red laser diodes (LDs) just above dorsal CA1 in both hemispheres (**Fig. 3A**).

**Figure 3.**
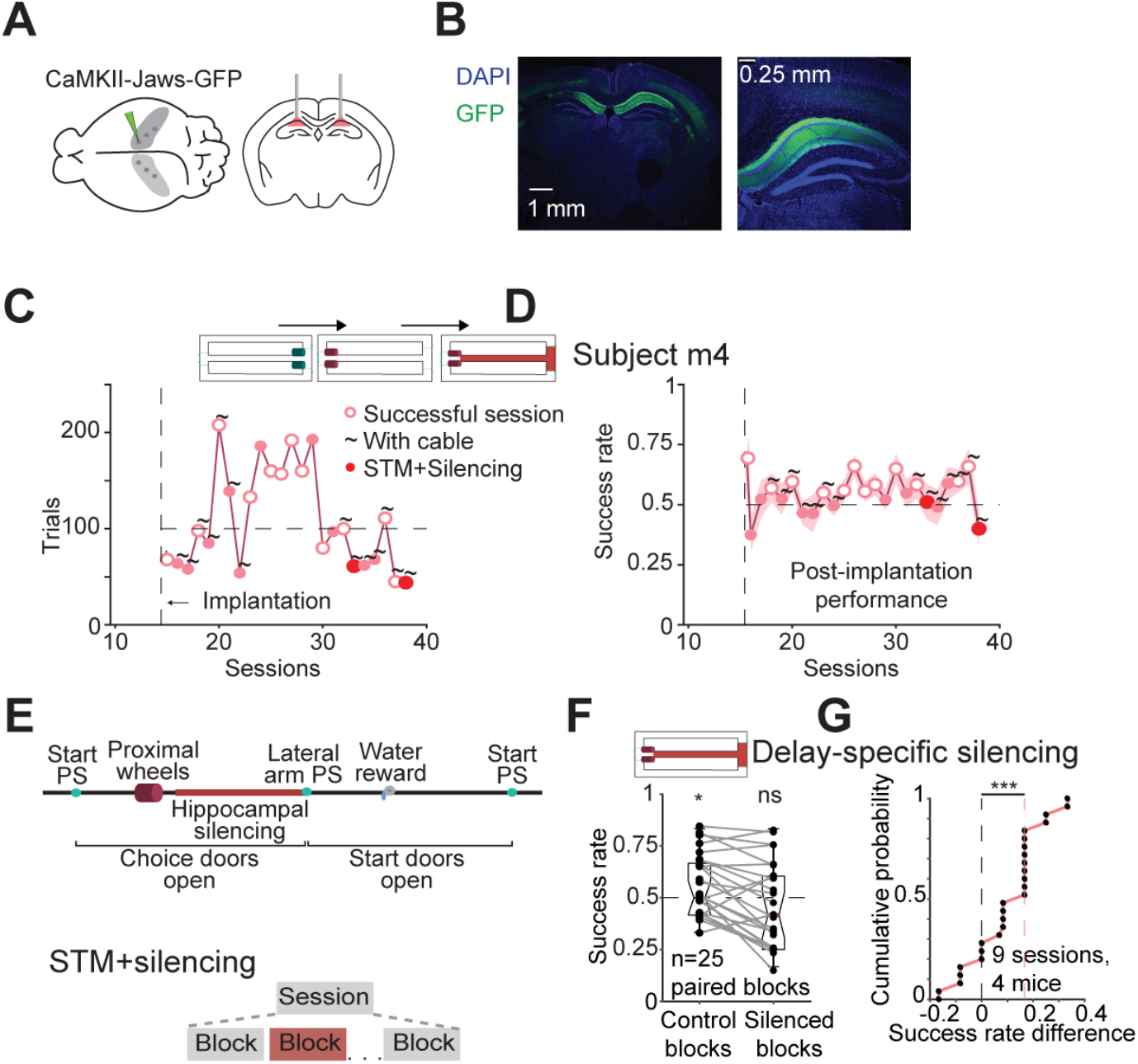
Bilateral silencing of dorsal CA1 during memory maintenance degrades success. (**A**) **Left**, Injection sites for CaMKII-Jaws-GFP viral vector. 150 nl were injected at every site, for a total of 20 sites in six craniotomies in two hemispheres. **Right**, Red LDs coupled to 200 μm optical fibers were implanted over dorsal CA1, one per hemisphere. (**B**) Wide-field coronal section of m4 (left; AP, −2 mm) and a closeup on the left hemisphere (right; AP, −1.6 mm). Jaws-GFP is expressed bilaterally in dorsal CA1. (**C**) Number of trials on the STM task by subject m4 after implantation. Horizontal line, number of trials criterion. Here and in **D**, all conventions are the same as in **Fig. 1DE**. (**D**) Success rates post-implantation. (**E**) **Top**, Timeline for a silenced trial on the STM task. **Bottom**, During STM+silencing sessions, blocks without and with silencing are interleaved. (**F**) Success rates during paired (consecutive) Control and Silenced blocks. ns/*: p>0.05/p<0.05, Wilcoxon’s test comparing to chance level, 0.5. (**G**) Silencing effect, defined as the success rate in every Control block minus the success rate in the following Silenced block. Dataset includes 50 blocks from nine sessions in four mice (**Table S3**). ***: p<0.001, Wilcoxon’s test.

Histological assessment showed that expression was confined to the dorsal hippocampus (**Fig. 3B, Fig. S3ABC**). Before commencing silencing sessions, we assessed post-implantation performance. After implantation, performance deteriorated in terms of number of trials per session (p=0.034, U-test; n=4) and success rate (p<0.001; **Fig. S2**). Nevertheless, 28/56 (50%) of the post-implantation unsilenced sessions were successful (p<0.001, Binomial test), with a median [IQR] success rate of 0.56 [0.50 0.62] (p=0.0014, Wilcoxon’s test; **Fig. S3E**). The number of trials per post-implantation session was 108 [85 155] (n=56 sessions, **Fig. S3D**). Thus, above chance performance was maintained after implantation.

To determine whether the tactile STM task is hippocampus dependent, we silenced the pyramidal cells of the dorsal hippocampus specifically during the memory phase of the task (**Fig. 3E**, top). Silencing was done in a closed-loop contingent on animal position^65^, confined to the maintenance and retrieval phases of the task. Namely, bilateral CA1 illumination was carried out from the moment the animal passed the proximal texture wheel, after sample encoding ended, until a choice was made, indicated by a lateral-arm photosensor (**Fig. 1A**). Furthermore, silencing was carried out during all trials of even-numbered blocks, whereas during the alternating Control blocks no silencing was applied (**Fig. 3E**, bottom).

We found that during Control blocks, performance was above chance level (n=25 blocks in 9 sessions; p=0.017, Wilcoxon’s test comparing to chance level, 0.5; **Fig. 3F**). In contrast, during the ensuing Silenced blocks, performance was not consistently different from chance (p=0.39; **Fig. 3F**). The delay-specific silencing induced a median performance decrease of 16.7% in success rate between consecutive blocks (p<0.001, Wilcoxon’s test; **Fig. 3G**). Similar silencing effects were observed for all individual subjects (range, [7.7%,16.7%]; n=4 mice; **Table 1**). Thus, success rate on the non-spatial STM task deteriorates following transient silencing of the bilateral dorsal hippocampus during the memory maintenance phase.

## Discussion

We developed a tactile STM paradigm for freely-moving mice which was learned by all mice within a dozen sessions. Transient silencing of the dorsal hippocampus during the maintenance epoch of the STM task decreased success, suggesting that intact CA1 activity is critical for the memory part of the task. To the best of our knowledge, this is the first time that a non-spatial STM task is being used by freely-moving mice, and the first direct evidence implicating the hippocampus in the maintenance of non-spatial STM in any species.

### Tactile STM task for freely-moving mice

Previous tactile STM tasks were developed only for head-fixed subjects^41,43–45^ but not for freely-moving mice (**Table S1**). In developing the paradigm, we hypothesized that passing by the textures would make the task easier to learn, compared with head-fixed tasks. For example, previous TD tasks for freely-moving rodents required 3-10 sessions to learn^55,66–69^ (**Table S2**), while similar TD tasks in head-fixed rodents required dozens-to-hundreds of sessions^49,50,52,54,70^. Indeed, all mice learned the present TD task within less than seven sessions (median, 5 sessions; **Table S3**). We also hypothesized that the organic nature of the task will motivate the subjects to perform many trials. In our TD task, mice ran more than a hundred trials per session, much higher compared with other TD tasks for freely-moving rodents (**Table S2**), where subjects perform up to a few dozen trials in a session^55,67–69^.

In the STM task, since the sampling of the texture was done *en passant*, delay duration was determined by track length and running speed. In designing the task, we hypothesized that ingraining the delay into the run may shorten learning duration compared with nose-poke or fixation delays. In previous sensory STM tasks, freely-moving rats^25,26,31,32,35,39,42^ and head-fixed mice^28,33,34,38,43^ required 5-90 sessions for learning the task (**Table S1**). However, learning the present STM task required only a dozen sessions (**Table S3**), and every single subject learned the task.

### Hippocampal involvement in STM

Previously, specific brain areas were assigned a causal role in STM by region-specific lesioning^10,12,71,72^ or silencing^36,37,39,41,44,45,73,74^ during maintenance and observing the behavioral effect. In all mice, we found that bilateral silencing of dorsal CA1 during memory maintenance degraded success in the tactile STM task. The decreased performance while CA1 is silenced specifically during the delay can be interpreted in at least three ways. One possibility is that animals memorized the stimulus and the delay-specific silencing interfered with the memory maintenance function. Then, the conclusion is that CA1 is necessary for the maintenance of sensory STM. An alternative is that subjects memorized correct motor responses rather than stimuli, drawing on the stimulus-response association already at the beginning of the middle arm. Then, the conclusion would be that the memory maintenance component of the task is largely spatial. A third possibility is that the animals prepared for a motor action at the T-junction. This possibility coincides temporally with motor memory and but corresponds to preparatory activity. There are of course “hybrid” options, wherein a subject remembers the stimulus at the beginning of the middle arm, draws on the stimulus-response association along the run, and then maintains the memory of the response. From the cognitive perspective, the first two options correspond to memory maintenance, and the distinction is equivalent to determining when the decision is made. Experimentally, the distinction could be made by silencing the hippocampus during distinct parts of the delay.

While STM corresponds to memory maintenance and is fully defined by duration^1^, WM has been segmented into sequential phases, thought to rely on different neuronal processes: encoding, maintenance, and retrieval^3^. Previous work concluded that silencing hippocampal input to the prefrontal cortex impairs encoding but not the maintenance of spatial WM^16^. Here, encoding occurred when the animals passed by the proximal wheels but hippocampal silencing began later, specifically during the maintenance and retrieval phases. The impairment of performance extends the previous findings, indicating that dorsal hippocampal activity is needed for WM maintenance and retrieval.

### Extensions and variations

All subjects learned the task within a dozen sessions, and therefore the paradigm provides a reliable manner to study tactile discrimination and delayed-response in freely-moving mice. The level of difficulty can be varied and a wide range of parameters can be explored. We used the most distinct grits on the texture wheels, P60 and P320. By using other combinations of textures, additional questions may be addressed. For instance, the just-noticeable difference of textures during a run can be studied using P60/P150, P60/P240, P60/P320, P150/P240, P150/P320, and P240/P320 pairs, yielding a psychometric curve. Delay duration could be controlled artificially by adding doors or a treadmill^13^ to the middle arm. A complementary application is to use both wheel pairs in the same trial, yielding a delayed comparison task^39,75^. The animal would sample one texture at the proximal wheels, run along the middle arm, and then sample the second texture at the distal wheels.

### Limitations

The presence of training trials effectively reduced the number of testing trials per session. The logic for including training trials was to facilitate rapid learning and minimize frustration. To evaluate the implications, m5 underwent one STM session that included only testing trials, where success rate was within the range of successful sessions (0.72, x-marked circle; **Fig. S2E**). Future work may evaluate whether training trials can be removed altogether for a well-trained mouse. The design of silencing experiments may also be changed: we chose to silence entire blocks to eliminate the possibility of error-leading-error bias. An alternative is to silence individual testing trials at random.

## Acknowledgements

We thank Yonatan Shapira and Shaked Zeierman for constructive comments. This work was supported by the Minducate Center (to S.S.); by the United States-Israel Binational Science Foundation grant 2015577 (to E.S.); by the European Research Council grant 679253 (to E.S.); by the Israel Science Foundation grant 638/16 (to E.S.); by the Israel Science Foundation FIRST Program grant 1871/17 (to E.S.); and by the Zimin Institute (to E.S.).

## Author contributions

E.S. conceived the project. S.S. and E.S. designed the apparatus and the experiments. S.S. and A.L. constructed the apparatus and implanted animals. S.S. and N.K. carried out experiments. S.S. and E.S. analyzed data. S.S. and E.S. wrote the manuscript, with input from all authors.

## Materials and Methods

### Experimental animals

Nine adult mice, two males and seven females, were used in this study (**Table 1**). Eight mice were hybrid (HYB), offspring of FVB/NJ females (JAX #001800, The Jackson Labs) and C57BL/6J males^76^. Compared to progenitors, hybrids exhibit reduced anxiety-like behavior, improved learning, and enhanced running behavior^76^. One mouse (m6) was a C57-derived offspring of an PV-Cre male (JAX #008069) and an Ai32 female (JAX #012569). After separation from the parents, animals were housed in groups of same-litter siblings until participation in experiments. Animals were held on a reverse dark/light cycle (dark phase, from 8 AM until 8 PM). In subjects m3 and m6, electrophysiological recordings and optical manipulations were carried out during some sessions. Recordings were carried out using a silicon diode-probe mounted on a micro-drive ^77^. Results of electrophysiological recordings are not included in the present report. All results were observed at the subject level (**Table 1**; **Table S3**; **Fig. S2**), and no differences were observed between subjects equipped or unequipped with diode-probes. All animal handling procedures were in accordance with Directive 2010/63/EU of the European Parliament, complied with Israeli Animal Welfare Law (1994), and approved by the Tel Aviv University Institutional Animal Care and Use Committee (IACUC #01-16-051, #01-20-049, and #01-21-052).

### Apparatus

The apparatus was a 140 x 50 cm figure-8 maze with two pairs of motorized texture wheels, located 120 cm apart at opposite ends of the middle arm (**Fig. 1A**). All sensors and actuators were controlled by a microcontroller (Arduino Mega) via custom designed electronic circuitry. The start area (L x W x H: 30 x 10 x 3 cm) was located at the beginning of the central arm (120 x 4 x 3 cm) and was connected to the end of the two lateral arms (140 x 10 x 3 cm). The passageways connecting the home box and the lateral arms were blocked by two transparent polycarbonate “start” doors. Two additional “choice” doors were located at the sides of the T-junction at the end of the central arm, blocking passage from the T-junction to the lateral arms. Every door was operated by a motor (DC 6V 30RPM Gear Motor, Uxcell) and was equipped with two limit switches (D2F-01L2, Omron). There were four photosensors (S51-PA-2-A00-NK, Datasensor), two in the start area, and one after each choice door. Water delivery was controlled by solenoid valves (003-0137-900, Parker). Every water port was connected to a different solenoid via flexible (ID, 1/16”, Tygon) tubing.

Every polycarbonate texture wheel (OD, 12 cm; width, 7 cm) was divided into five facets. Four facets were coated with sandpaper of different grits: P60 (coarse, 269 μm particle diameter), P150 (100 μm), P240 (58.5 μm), and P320 (extra fine, 46.2 μm). The fifth (null) facet was plain polycarbonate. Tactile stimulation was given by rotating the wheels using 12V/350 mA stepper motors (200 steps/rev, NEMA-17, Adafruit). By default, the wheels were set with the null facet facing the central arm. White LEDs (c512A-WNS-CZ0B0152, Cree) were installed in each wheel to emphasize the active wheels, and were lit when the wheels were in use.

### Water deprivation protocol

Mice were trained on a TD task and/or on a tactile delayed-response (STM) task. Every session was conducted on a different day. At the beginning of the training period, animals were housed one per cage and placed on a water-restriction schedule that guaranteed at least 40 ml/kg of water every day, corresponding to 1 ml for a 25 g mouse. Training was carried out five days a week, and animals received free water on the sixth day. Reward volume differed between mice and sessions, ranging [4,20] μl. The exact volume was determined by the experimenter before each session based on familiarity with the specific animal. Mice received a water reward after every training trial and a 30-50% larger reward after every successful testing trial.

### Light sources and surgery

Implants comprised of two red (638 nm) laser diodes (LDs; HL63603TG, Ushio) coupled to 2 cm long 200 μm diameter optical fibers (FG200UEA, Thorlabs). Diodes were driven by a precision multi-channel current source^77^. The maximal driving current used was 50 mA, resulting in light power of 3.6±1.1 mW measured at the tip of the fiber (mean±SD over n=8 light sources in four mice).

Four hybrid mice (m4, m7, m8, and m9) were injected bilaterally with CaMKII-Jaws (rAAV5/CaMKII-Jaws-KGC-GFP-ER2; 5.2 x 10^12^ IU/mL; University of North Carolina viral core facility, courtesy of E.S. Boyden) to express Jaws in PYRs. The viral vector was injected into the dorsal hippocampus at each of 20 sites (AP −1.23, ML ±0.75, DV 1.3, 1.5 and 1.7; AP −1.6, ML ±1.1, DV 1, 1.2 and 1.4; and AP −2, ML ±1.75, DV 1.05, 1.25, 1.45 and 1.65; 150 nl/site) for a total of 1.5 μl per hemisphere. Following the injections, two diode-coupled optical fibers were implanted, one in the central penetration site of every hemisphere (AP −1.6 mm, ML ±1.1 mm, DV 1.0 mm) under isoflurane (1%) anesthesia ^77^.

### Tactile discrimination task

Five mice (m1, m2, m3, m4, m5) were trained on the TD task. In the TD task, the textures were presented at the end of the middle arm using the internally-illuminated distal wheels, next to the choice T-junction. In all sessions, P60 (coarse) textures were associated with leftward runs, and P320 (extra fine) textures were associated with rightward runs (**Fig. S1A**). Allocation of texture cues to trials was pseudorandom, designed to contradict strategies the mouse can employ. To prevent subjects from developing automated strategies that allow obtaining reward without learning the task, the strategy that the mouse is most likely to be taking was estimated online^78^. Thus, the algorithm estimated the most probable instantaneous strategy and provided trials that minimize the effectiveness of the strategy.

Initially, each mouse was acquainted with the task in a set of shaping sessions (median [range]: 4 [4,6] sessions; n=5 mice; **Fig. S1B**; **Table S3**). Shaping sessions included only training trials. In the training trials, only the correct choice door opened following the presentation of a texture stimulus, forcing the animal to choose the rewarded side. Mice had to reach a criterion of 100 trials per session before moving on to learning sessions. In every session, the mice were free to perform the task until losing interest, identified by prolonged periods of rest and attempts to climb the walls.

Post-shaping sessions were divided into blocks, and every block included four training trials and 12 testing trials (**Fig. 1B**). A single testing trial proceeded as follows: (1) Run: Once the animal passed a start photosensor, the start doors closed and the choice doors opened (**Fig. 1C**). (2) Stimulus: Upon arrival at the distal wheels, the stimulus could be sampled. (3) Response: The animal chose a direction at the T-junction and went through one of the two open choice doors. Once the animal passed a lateral arm photosensor, the choice doors closed and the start doors opened. (4) Reward: If the animal made a correct choice, a water reward was immediately available at the corresponding water port. Because the choice doors were already closed, the animal could not go back to the T-junction but was free to consume the reward and return to the start area.

### Tactile STM task

Seven mice (m2, m4, m5, m6, m7, m8, m9) were trained on the tactile delayed-response (STM) task. In the STM task, textures were presented at the beginning of the middle arm using the internally-illuminated proximal wheels. The distal wheels were kept stationary, with the indicator LEDs off and the null facets facing the middle arm. The stimulus-response association during the STM task was the same as in the TD task, namely textures P60 and P320 corresponded to left and rightward runs, respectively (**Fig. S1A**). The presentation of the textures at the beginning of the middle arm requires memory maintenance throughout the middle arm run. The median [IQR] delay duration, estimated based on 710 trials in seven sessions of subject m9, was 3.3 [2.7 4.3] s. Training on the STM task began with shaping sessions (median [range]: 2 [1,4] sessions; n=7 mice; **Table S3**). As in the TD task, the criterion for moving from shaping to learning sessions was 100 trials per shaping session. All other procedures were identical to those described for the TD task.

### Optogenetic manipulations

Implanted animals (n=4 mice; **Table 1**) were equipped with a three-axis accelerometer (ADXL-335, Analog Devices) for monitoring head movements. Head position and orientation were tracked using two head mounted LEDs, a machine vision camera (ace 1300-1200uc, Basler), and a dedicated system, “Spotter”. Specifically, head position was calculated by the mean position of the two head mounted LEDs, and head orientation was estimated by the normal to the line that connects the two LEDs. Stability and continuity were provided by a rigid body Kalman filter realized in real-time^65^. Mouse behavior was visualized online by the experimenter, and registered automatically by photosensor crossing times. During every silencing session animal kinematics, the currents applied to the LDs, and all digital events were recorded by an RHD2000 evaluation board (Intan Technologies) at 20 kHz (16 bits).

During silencing sessions, a virtual position sensor was created right after the proximal pair of wheels using Spotter, generating a digital pulse that was routed to a digital signal processor (DSP; RX8, Tucker Davis Technologies). The DSP also received input from the lateral arm photosensors. Illumination was applied from the moment that the subject crossed the virtual sensor until a lateral arm photosensor was crossed, corresponding to the maintenance and retrieval phases. Closed-loop illumination was carried out on every training and testing trial in alternate (even) Silenced blocks. No silencing occurred during the first block. During the Silenced blocks, the DSP generated a voltage command, given to the precision multi-channel current source that drove the implanted LDs to generate red light.

### Training on the STM task

Three mice (m2, m4, and m5) were first trained on the TD task and subsequently on the STM task (**Fig. S1C**, top, “Pretrained”). Shaping sessions for the STM task lasted a single session for all pretrained mice (**Table S3**), presumably reflecting the fact that the animals were already acquainted with the apparatus and the general rules. Indeed, the STM shaping phase included 161, 132, and 190 trials for mice m2, m4, and m5, respectively. For the same mice, the ensuing STM learning phase required a total of 576, 194, and 828 STM trials (**Fig. S1K**) that spanned 3, 1, and 4 sessions (**Table S3**; **Fig. S1L**). After learning, the three mice carried out a total of 33 sessions, of which 28 (85%) were successful (**Fig. S2BDE**).

Notably, success rates while performing the STM task (median, 0.63; n=100 performance sessions; **Fig. 2E**) were lower than success rates during the TD task (0.83; n=15 sessions; p<0.001, U-test; **Fig. 1G**; **Table 1**). Among the mice that performed both tasks with identical stimuli, success rates were lower in the STM task (median, 0.62, n=32 sessions) compared with the TD task (0.88, n=7; p<0.001, U-test, **Fig. S1J**). Comparing success rates of only the mice naïve to each task (i.e., mice that learned each task directly: m1-m5 on the TD task, and m6-m9 on the STM task) yielded similar results (TD; median, 0.83, n=15 sessions; STM: 0.64, n=68; p<0.001, U-test).

Learning the TD task first can benefit learning the STM task via generalization^79^, or detriment learning of the STM task due to extinction or interference ^80^. To determine whether direct training on the STM is possible and if so beneficial, we trained four other mice (m6, m7, m8, m9) directly on the STM task (“naïve” mice). The shaping phase of the naïve mice lasted 4, 2, 4, and 4 sessions (**Table S3**), and included 255, 115, 219 and 223 trials, not consistently different from the shaping phase of the five TD mice (range, [3,6] sessions; p=0.4, U-test).

Akin to the pretrained mice, the naïve subjects went through a learning phase. Due to a technical issue, two of the naïve mice (m8, m9) were trained on the STM in the light (**Fig. S2HI**, light pink background). The technicality proved insightful. The learning phase lasted 19 and 20 sessions for the two mice (**Table S3**; **Fig. S1L**, open circles), considerably longer compared with the learning phase of the dark-trained naïve mice. After 19 and 18 sessions in the light, the two light-trained naïve animals were transitioned to the standard (dark) training conditions, and achieved learning within up to two sessions (**Table S3**). In sum, the learning phase of the naïve mice lasted 5, 7, 19, and 20 sessions. Thus, a dozen sessions are required to learn the STM (**Table S3**), either following the TD task (12, 9, and 13; **Fig. S1L**) or directly (9 and 9; **Fig. S1L**).

The pretrained mice (n=3; **Fig. S2BDE**) experienced four phases prior to STM performance: TD shaping and performance (**Fig. S1M**, green hues), followed by STM shaping and learning (**Fig. S1M**, pink hues). In contrast, the naïve mice (n=4; **Fig. S2FGHI**) experienced only two phases of training prior to STM performance: STM shaping and learning (**Fig. S1C**). The two cohorts did not exhibit consistent differences in the number of trials required for learning (p=0.23, for all mice; p=0.8 when excluding m8 and m9, U-test; **Fig. S1K**), or in the number of learning sessions on the task (p=0.06, for all mice [pretrained, n=3; naïve, n=4]; p=0.2 when excluding m8 and m9; U-test; **Fig. S1L**). All performance sessions of the naïve mice were carried out in the dark, with a total of 59/68 (87%) successful sessions (**Fig. S2FGHI**). The success rates during the performance phase were not consistently different between the pretrained and naïve mice (p=0.2, U-test; **Fig. 2E**). However, during the performance phase, pretrained mice performed more trials per session (p=0.005, U-test; **Fig. 2D**). Thus, even under suboptimal training conditions, all mice successfully learn the STM task.

### Effect of the chronological day of training

During the performance phase, some fluctuations were seen in the success rates, in both tasks (**Fig. S2**). Reporting session performance sequentially while disregarding the actual calendar days (e.g., Sunday, Monday, Tuesday) may obscure fluctuations due to the session ordinate in a week. For example, success rate may depend on whether the session was carried out on the first or the last day of training during a week. We found that the number of trials depended on the chronological day of the training, in both the TD task and the STM task (**Fig. S1EG**), but success rate did not (**Fig. S1FH**). Specifically, the number of trials depended on the chronological day of the training (rank correlation coefficient, 0.33; p <0.001, permutation test; **Fig. S1G**). On the first day after a break, mice performed a median [IQR] of 104 [70 136] trials (n=30 STM sessions), whereas on the third day the same mice performed 150 [118 177] trials (n=20 sessions). Both chronological day and the number of trials contributed to the variability of the success rate (R^2^, 0.044; p<0.001, Wilcoxon signed-rank test; **Fig. S1I**). However, success rate in the STM task did depend on the combination of chronological day and the number of trials (**Fig. S1I**). The chronological day of training may influence performance in at least two ways: the motivation of the animal which may be embodied by the number of trials; and the training momentum, realized as across-session long-term learning.

### Quantification of behavior

Sessions with more than 40 trials (training and testing) were considered valid. The criterion for moving from the shaping phase to the learning phase was performing 100 trials in a single session. The criterion for moving from the learning phase to the performance phase was two valid successful consecutive sessions. Performance was considered successful whenever p<0.05 in a Binomial test comparing to chance level, 0.5.

### Histological analyses

After conclusion of all experiments, fiber implanted mice were deeply anesthetized with pentobarbital (100 mg/kg) and perfused via the left ventricle with 0.1 M phosphate buffered saline (PBS; pH 7.4) followed by 4% paraformaldehyde (PFA). Brains were removed and postfixed overnight in PFA and then transferred to PBS. Coronal sections (50-70 μm) were cut (VT1000S, Leica), collected in PBS, and mounted in Fluoromount with DAPI (F6057-20ML, Sigma). Sections were imaged using a fluorescence microscope (Axio Scope A1, Zeiss) or a confocal laser scanning microscope (SP8, Leica).

### Statistical analyses

In all statistical tests a significance threshold of α=0.05 was used. All descriptive statistics (n, median, IQR, range, SEM) can be found in the text, figures, figure legends, and tables. Differences between medians of two unpaired groups were tested with Mann-Whitney’s U-test (two tailed). Wilcoxon’s signed-rank test was employed to determine whether the medians of two paired groups are distinct (two-tailed) and whether a group median is distinct from zero (two tailed). To estimate whether a given fraction was larger than expected by chance, an exact Binomial test was used (one tailed). Permutation tests were used to estimate significance of rank correlation coefficients. In all figures, ns: p>0.05; *: p<0.05; **: p<0.01; ***: p<0.001.

## Supplementary Materials

**Figure S1.**
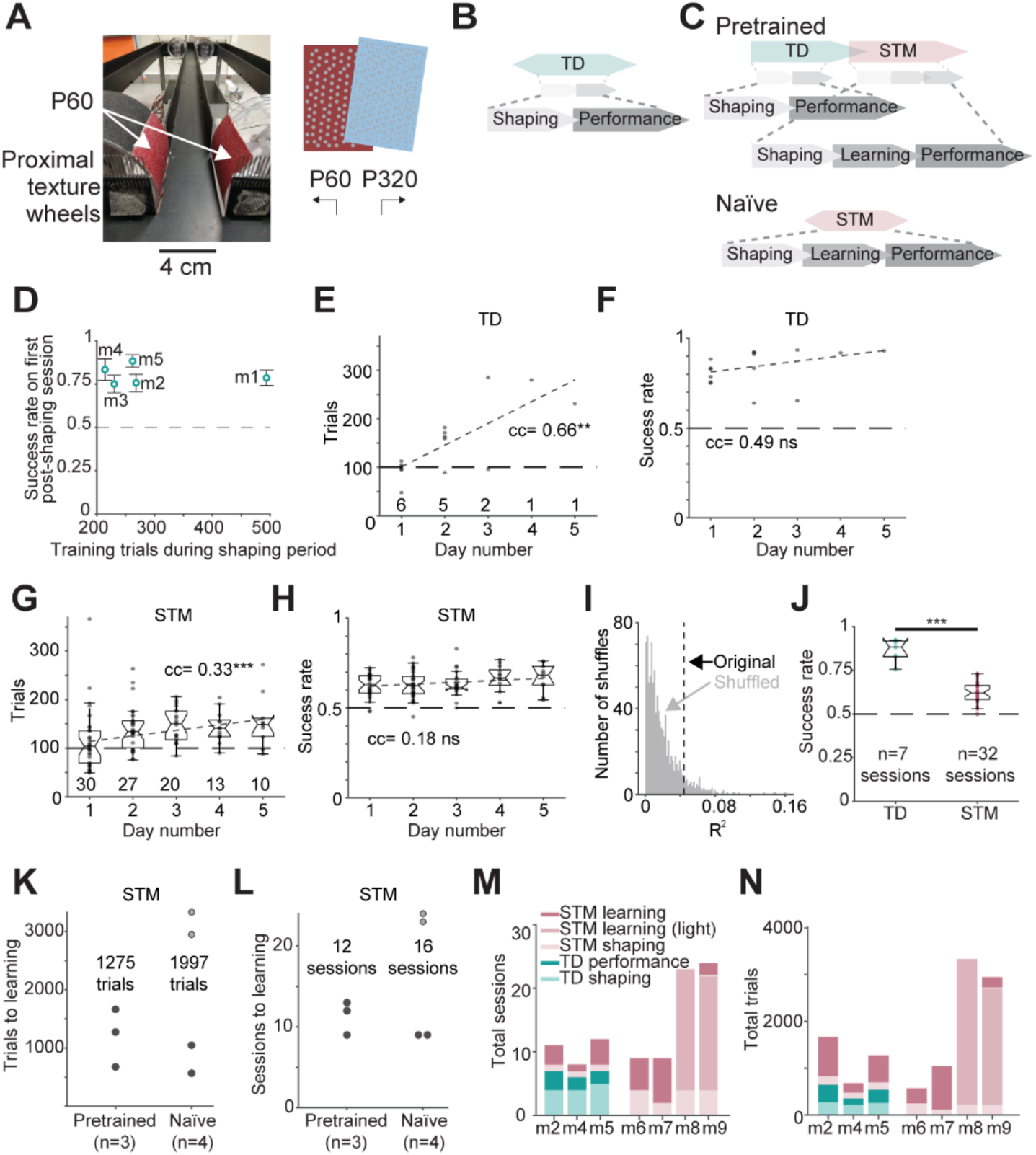
Training procedure and dependency of performance on the chronological training day. (**A**) **Left**, A view of the proximal wheels and the middle arm from the perspective of a mouse at the start area. **Right**, Stimulus-response contingency. Right and left turns are associated with fine and coarse tactile stimuli, respectively. (**B**) Procedure used to train mice on the TD task. Subjects begin every task with a shaping phase until performing a hundred training trials per session. Shaped mice undergo a learning phase that ends at the first of two consecutive successful sessions. (**C**) Two approaches used to train mice on the STM task. **Top**, Pretraining on the TD task before the STM task. **Bottom**, Direct training of naïve mice on the STM task. (**D**) Mice learn the TD task during the shaping phase, performing successfully from the very first post-shaping session. The number of shaping sessions is 4, 4, 6, 4, and 5, for mice m1-m5. Error bars, SEM. (**E**) The number of trials depends on the chronological day number during performance sessions of the TD task. Day number 1 is the first day after a break (e.g., a weekend). Here and in **FGH**, ns/**/***: p>0.05/p<0.01/p<0.001, permutation test; dashed line, robust fit. The number of sessions per day is denoted below every dot plot. (**F**) The success rate does not depend on the chronological day during TD performance sessions. (**G**) Dependence of the number of trials on the chronological day during STM performance sessions. Here and in **H**, box plot conventions are identical to **Fig. 2D**. (**H**) Dependence of success rate on chronological day during STM performance sessions. (**I**) Both chronological day and the number of trials contribute to success rate variability in the STM task. Median R^2^ with shuffled labels, 0.015. (**J**) Among the three mice that performed both tasks with identical stimuli, success rates were lower during the STM task. ***: p<0.001, U-test. (**K**) Total number of trials required for STM learning in every subject. Subjects m8 and m9 were initially trained in the light, denoted here and in **L** by empty circles. (**L**) Total number of sessions required for STM learning for every subject. (**M**) Number of sessions during each phase for every mouse learning the STM task. The median [range] number of total sessions required for learning the STM task in the dark is 9 [9,13] (n=5 mice). (**N**) Number of trials during each phase for every mouse learning the STM task. The median [range] number of total trials required for learning the STM task in the dark is 1049 [645,1664] (n=5 mice).

**Figure S2.**
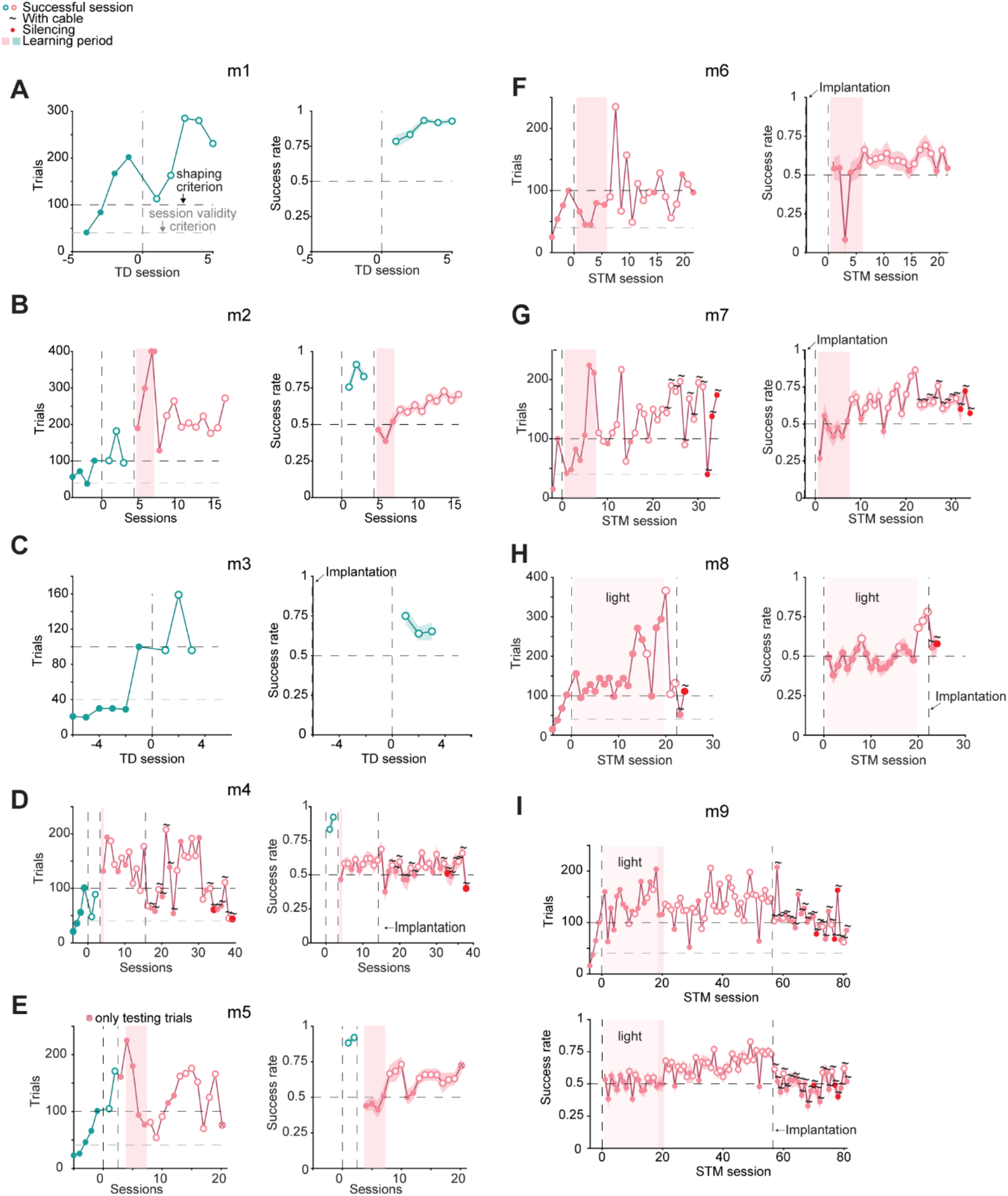
Learning curves of every mouse. (**A-I**) **Left**, Number of trials performed during the shaping and performance phases. Horizontal black line indicates the 100-trials shaping criterion, and the horizontal gray line indicates the 40-trials session validity criterion. **Right**, Success rates during every session in the learning and performance phases. Horizontal line indicates chance level, 0.5, and the vertical dashed line at 0 marks the transition between shaping and learning or performance phases. Open circles denote successful sessions (p<0.05 in a Binomial test, comparing to chance level). Error band, SEM. (**C**) In subject m3, the implantation took place prior to initiation of training, marked here and in **FG** by a second dashed vertical line (here, at −6). (**D**) In subject m4 the implantation of the LD-coupled optical fibers is indicated by a vertical dashed line. (**H-I**) Number of trials performed during every session in the learning and performance phases of m8 and m9. Lighter pink background indicate training in light. Other conventions are the same as in **D**.

**Figure S3.**
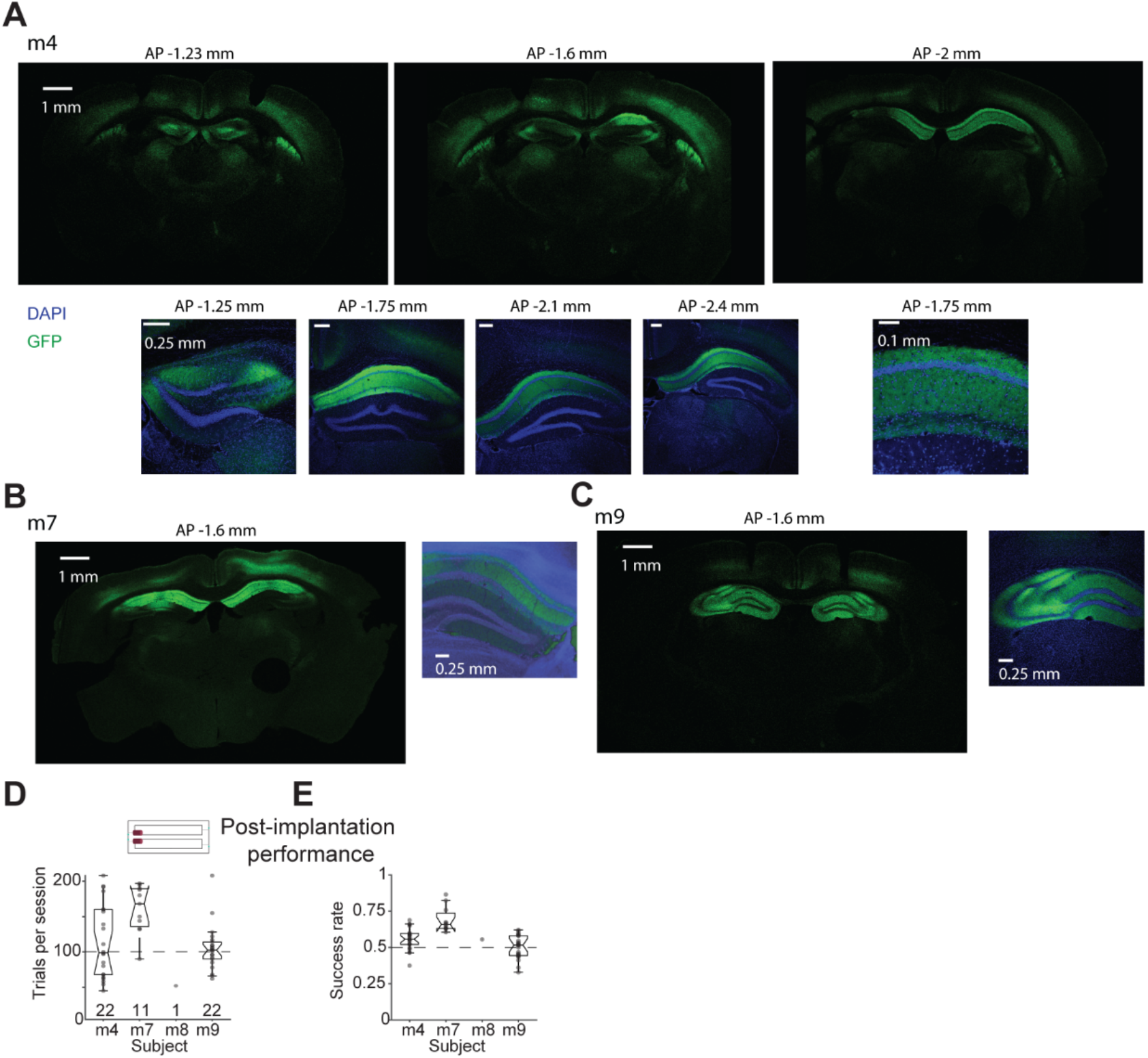
Histological validation of Jaws-GFP expression in dorsal CA1 and post-implantation performance. (**A**) Jaws-GFP is expressed in dorsal CA1. **Top**, Wide-field coronal sections along the anterior-posterior (AP) axis of m4. Here and in **B-C**, green, green fluorescent protein (GFP) fluorescence; blue, 4’,6-diamidino-2-phenylindole (DAPI) fluorescence. **Bottom**, Magnified views of the left hippocampus along the anterior-posterior axis. (**B**) **Left**, Wide-field coronal section of m7. **Right**, Magnified view of the right hippocampus. (**C**) **Left**, Wide-field coronal section of m9. **Right**, Magnified view of the right hippocampus. (**D**) Success rates and number of trials on the STM task after implantation. Silencing sessions are not included. Here and in **E**, Box plot conventions are the same as in **Fig. 2D**. The number of sessions per mouse is denoted below every box plot. (**E**) Success rates on the STM task after implantation.

### Supplementary tables

**Table S1.** STM tasks.

| Task type | Modality | Species | Preparation | Subjects trained | Subjects that learned | Training duration<sup>1</sup> | Trials per session<sup>1</sup> | Success rate<sup>1</sup> | Year of publication | Source |
| --- | --- | --- | --- | --- | --- | --- | --- | --- | --- | --- |
| MWM | spatial | rats | FM | N/A | 31 | 10 trials | [4,8] | <8 s latency | 1982 | (Morris *et al.*, 1982) |
| MWM | spatial | rats | FM | N/A | 16 | 4 sessions | 1 or 4 | [10,30] s latency | 2014 | (Plescia *et al.*, 2014) |
| DNMP | spatial | mice | FM | N/A | 29 healthy<br>21 HD | 8 days | 80 [75,90] | [0.7,0.8] | 2016 | (Yhnell *et al.*, 2016) |
| DNMP | spatial | rats | FM | N/A | 18 | N/A | 28 [20,36] | 0.8 | 2023 | (George *et al.*, 2023) |
| 2AFC | spatial | rats | FM | N/A | 5 | 3 days | 29±2.6 | 0.93<br>[0.89,0.96] | 2003 | (Baeg *et al.*, 2003) |
| 2AFC | spatial | rats | FM | N/A | 20 | 11.5 [9,13] days | 40 | 0.83<br>[0.8,0.87] | 2007 | (Ainge *et al.*, 2007) |
| 2AFC | spatial | rats | FM | N/A | 15 | >6 days | 40 | 0.75 | 2008 | (Yoon *et al.*, 2008) |
| 2AFC | spatial | rats | FM | N/A | 6 | 3-15 sessions | 40 | 0.75 | 2013 | <sup>13</sup> |
| 2AFC | spatial | mice | FM | N/A | 18 | 4 sessions | 10 | 0.85±0.05 | 2014 | (Pioli *et al.*, 2014) |
| 2AFC | spatial | mice | FM | 27 | 27 | >5 sessions | N/A | 0.77<br>[0.72,0.82] | 2015 | <sup>16</sup> |
| 2AFC | spatial | rats | FM | N/A | 4 | 65 days | 107<br>[94,109] | 0.87<br>[0.77,0.9] mean<br>[range] | 2019 | (Holleman *et al.*, 2019) |
| DNMS | olfactory | rats | FM | N/A | 3 | N/A | 200 | 0.95 | 1993 | (Lu *et al.*, 1993) |
| DNMS | olfactory | rats | FM | >4 | 4 | 20 days | N/A | 0.92 | 2008 | (Fujisawa *et al.*, 2008) |
| DNMS | olfactory | mice | HF | 253 | 12 | 5-7 days | [100,200] | 0.95 | 2014 | (Liu *et al.*, 2014) |
| DD and DC | olfactory | mice | HF | N/A | 12 | 7-8 days | 100±110 | 0.8 | 2023 | (Reuschenbach *et al.*, 2023) |
| CNM | auditory | rats | FM | >8 | 8 | [9,41] sessions | 120 | 0.8 | 1987 | (Sakurai, 1987) |
| DD | auditory | mice | HF | N/A | 9 | several weeks | >50 | 0.84 | 2017 | (Kamigaki & Dan, 2017) |
| DC | auditory | rats | FM | N/A | 25 | 90 sessions | 200 | 0.78<br>[0.59,0.93] | 2018 | (Akrami *et al.*, 2018) |
| DD | visual | mice | HF | N/A | >=6 | 34 [26,44] sessions | 149<br>[126,162] | 0.83<br>[0.74,0.92] | 2012 | (Harvey *et al.*, 2012) |
| DD | visual | rats | FM | N/A | 5 | 55 days | 96 | [0.7,0.82] | 1980 | <sup>25</sup> |
| DD | visual | rats | FM | 16 | 12 | 26-36 weeks | 256 | [0.75,0.8] | 1992 | <sup>26</sup> |
| DC | tactile | rats | FM | N/A | 11 | [44,71] sessions | [200,400] | 0.71<br>[0.67,0.8] | 2014 | (Fassihi *et al.*, 2014) |
| DD | tactile | mice | HF | N/A | 22 | 3 weeks | 422±115 | 0.69±0.05 | 2014 | (Guo *et al.*, 2014) |
| DD | tactile | mice | FM | 7 | 7 | 12 [9,23] sessions | 132 [98,157] | 0.63 [0.59,0.84] | 2023 | **Present study** |

<sup>1</sup> Median [range] or mean±SD, unless stated otherwise.
Abbreviations: 2AFC, two-alternative forced choice; CNM, continuous non-matching-to-sample; DC, delayed comparison; DD, delayed discrimination; DMS, delayed match to sample; DNMP, delayed non-match to position; DNMS, delayed non-match to sample; FM, freely moving; GNG, Go/No-Go; HF, head fixed; MWM, Morris water maze.

**Table S2.** Tactile discrimination tasks.

| Task type | Species | Preparation | Subjects trained | Subjects learned | Training duration [sessions]<sup>5</sup> | Trials per session<sup>5</sup> | Success rate<sup>5</sup> | Year of publication | Source |
| --- | --- | --- | --- | --- | --- | --- | --- | --- | --- |
| OR | rats | HF | 34 | 26 | 11 [3,23] | 63±19 | 0.8 | 2006 | (Knutsen *et al.*, 2006) |
| OR | mice | FM | 16 | 12 | 3 days | 5 min of exploring | 62%<sup>6</sup> | 2013 | (Wu *et al.*, 2013) |
| GNG | mice | HF | N/A | 19 | 9 [7,14] | 300 [150,660] | 0.90 ± 0.003 | 2010 | (O'Connor *et al.*, 2010) |
| TD<sup>1</sup><br>GNG | mice | HF | N/A | 10 DD<br>5 GNG | TD 118 [93,157]<br>GNG 94.8 [24,147] | 161 (mean) | TD 0.75±0.05<br>GNG 0.8 [0.78,0.98] | 2021 | (Rodgers *et al.*, 2021) |
| TD<sup>2</sup> | rats | FM | N/A | 7 | 17 [15,28] | 20 | 0.85-1 | 1989 | (Guić-Robles *et al.*, 1989) |
| TD<sup>2</sup> | mice | FM | N/A | 20 | 3 | 36 | 0.9 | 1991 | (Lipp & Van der Loos, 1991) |
| TD<sup>2</sup> | rats | FM | N/A | 9 | 17 [15,28] | 20 | [0.85,1] | 1992 | (Guić-Robles *et al.*, 1992) |
| TD<sup>2</sup> | rats | FM | >8 | 8 | N/A | 35 [28,40] | 0.75 | 2007 | (von Heimendahl *et al.*, 2007) |
| TD<sup>3</sup> | rats | FM | N/A | 22 | [6,9] | >50 | 0.75 | 2001 | (Krupa *et al.*, 2001) |
| TD<sup>4</sup> | rats | HF | 14 | 11 | [61,225] sessions | [50,100] | Stable performance<sup>7</sup> | 2007 | (Mehta *et al.*, 2007) |
| TD | rats | FM | N/A | 7 | N/A | 108 | [0.7,0.95] | 2016 | (Grion *et al.*, 2016) |
| 2AFC | mice | HF | N/A | 3 | 17 [17,26] | 230 [200,380] | [0.8,0.95] | 2013 | (Mayrhofer *et al.*, 2013) |
| 2AFC | mice | FM | 5 | 5 | 5 [5,7] | 113 [95,182] | 0.83 [0.73,0.88] | 2023 | **Present study** |

<sup>1</sup> Shape-based TD.<sup>2</sup> Texture-based TD.<sup>3</sup> Aperture-based TD.<sup>4</sup> Angle-based TD.<sup>5</sup> median [range] or mean±SD, unless stated otherwise.<sup>6</sup> The measure of success was the time spent investigating the novel texture divided by time spent investigating the familiar or novel textures during the testing phase, minus 1.<sup>7</sup> Statistically significant differences in rewarded and unrewarded responses over multiple sessions.
Abbreviations: 2AFC, two-alternative forced choice; FM, freely moving; GNG, Go/No-Go; HF, head fixed; OR, object recognition; TD, tactile discrimination.

**Table S3.** Number of sessions in every task.

| Animal ID | TD shaping | TD learning | Total to TD | TD performance | STM shaping | STM learning | Total to STM | STM performance | Silencing |
| --- | --- | --- | --- | --- | --- | --- | --- | --- | --- |
| m1 | 4 | 1 | 5 | 5 | - | - | - | - | - |
| m2 | 4 | 1 | 5 | 3 | 1 | 3 | 12 | 9 | - |
| m3 | 6 | 1 | 7 | 3 | - | - | - | - | - |
| m4 | 4 | 1 | 5 | 2 | 1 | 1 | 9 | 10 | 2 |
| m5 | 5 | 1 | 6 | 2 | 1 | 4 | 13 | 13 | - |
| m6 | - | - | - | - | 4 | 5 | 9 | 16 | - |
| m7 | - | - | - | - | 2 | 7 | 9 | 13 | 3 |
| m8 | - | - | - | - | 4 | 19 (0)<sup>1</sup> | 23 | 3 | 1 |
| m9 | - | - | - | - | 4 | 20 (2)<sup>1</sup> | 24 | 36 | 3 |
| **Median** | **4** | **1** | **5** | **3** | **2** | **5** | **12** | **13** | **2.5** |

<sup>1</sup> Total number of STM learning sessions (number of STM learning sessions carried out in the dark).

